# Employing single-stranded DNA donors for the high-throughput production of conditional knockout alleles in mice

**DOI:** 10.1101/195651

**Authors:** Denise G. Lanza, Angelina Gaspero, Isabel Lorenzo, Lan Liao, Ping Zheng, Ying Wang, Yu Deng, Chonghui Cheng, Chuansheng Zhang, Matthew N. Rasband, John R. Seavitt, Francisco J. DeMayo, Jianming Xu, Mary E. Dickinson, Arthur L. Beaudet, Jason D. Heaney

## Abstract

The International Mouse Phenotyping Consortium is generating null allele mice for every protein-coding gene in the genome and characterizing these mice to identify gene-phenotype associations. To test the feasibility of using CRISPR/Cas9 gene editing to generate conditional knockout mice for this large-scale resource, we employed Cas9-mediated homology driven repair (HDR) with short and long single-stranded oligodeoxynucleotides (ssODNs and lssODNs). Using pairs of guides and ssODNs donating loxP sites, we obtained putative conditional allele founder mice, harboring both loxP sites, for 23 of 30 genes targeted. LoxP sites integrated *in cis* in at least one F0 for 18 of 23 targeted genes. However, loxP sites were mutagenized in 4 of 18 *in cis* lines. HDR efficiency correlated with Cas9 cutting efficiency but was not influenced by ssODN homology arm symmetry. By contrast, using pairs of guides and a single lssODN to introduce a loxP-flanked exon, conditional allele founders were generated for all 4 genes targeted. Our studies demonstrate that Cas9-mediated HDR with pairs of ssODNs can generate conditional null alleles at many loci, but reveal inefficiencies when applied at scale. In contrast, lssODNs are amenable to high-throughput production of conditional alleles when they can be employed.

## INTRODUCTION

Over the last decade the ease with which mammalian genomes can be modified *in vitro* and *in vivo* has substantially increased with the advent of gene editing technologies. Protein-guided nucleases (e.g. zinc finger nucleases and TALENS) and RNA-guided endonucleases (e.g. CRISPR/Cas9) enable genome editing at specific targeted sites in genomes through the generation of double-strand breaks and the subsequent repair of the double-strand breaks (DSBs) through imprecise non-homologous end joining (NHEJ) or precise homology-directed repair (HDR) with an exogenous DNA donor template (Sander and Joung 2014). Due to its ability to direct DSBs to precise locations in the genome with high efficiency (Gaj et al. 2013), the ease of reagent design, and the relative low cost of reagent production, the CRISPR/Cas9 system has become the preferred technology for germline gene editing in dozens of mammalian species, including mouse (Carroll 2014, Doudna and Charpentier 2014, Liu et al. 2017). Numerous publications have illustrated the ease of generating novel mouse lines from injections of Cas9 mRNA or protein and synthetic single guide RNAs (sgRNAs) (Wang et al. 2013, Wu et al. 2013, Yang et al. 2013, Horii et al. 2014, Chu et al. 2016, Andersson-Rolf et al. 2017, Chen et al. 2017, Ma et al. 2017) and reported methods to facilitate the production of edited alleles (Yoshimi et al. 2014, Miura et al. 2015, Boroviak et al. 2016, Ma et al. 2017).

While NHEJ-mediated null allele production in mice is highly efficient (Yang et al. 2013, Singh et al. 2015), conditional allele generation has proven to be more difficult. Adapting traditional plasmid-based methods to target conditional alleles into the genome of mouse embryos has been attempted but has proven to be inefficient (Wang et al. 2013, Yang et al. 2013). Furthermore, production of plasmid donors is time consuming and modifying existing plasmid-based gene targeting constructs from existing IMPC resources for CRISPR/Cas9-mediated HDR would be time and cost prohibitive for high-throughput production (Jung et al. 2017). As an alternative, short (≤200 bases) single-stranded oligodeoxynucleotide donors (ssODNs) are fast and relatively inexpensive to produce. Generation of conditional null alleles requires a pair of single guide RNAs (sgRNAs) targeting intronic sequences flanking exon(s) and a pair of loxP-containing ssODNs; this approach requires two DSBs and two independent HDR events to generate a conditional allele. The reported conditional founder rates per gene attempted has been less than or equal to 20% using pairs of sgRNAs and ssODNs (Yang et al. 2013, Lee and Lloyd 2014, Bishop et al. 2016, Miano et al. 2016, Ma et al. 2017). Importantly, each previous study using ssODNs to create conditional null alleles in mice targeted a single gene or a small subset of genes each under different conditions. Long single-stranded oligodeoxynucleotide donors (lssODNs) are also relatively easy and cost effective to produce, appear able to proceed with a single HDR event, and have recently been shown to be highly efficient at generating both conditional and reporter knock-in (KI) alleles (Miura et al. 2015, Quadros et al. 2017). Thus, ssODNs and lssODNs appear to be the most cost effect and efficient methods of conditional allele production in mice. Whether the apparent efficiencies of single-stranded donor DNAs will be maintained when systematically applied at multiple loci and at scale has not been evaluated.

The International Mouse Phenotyping Consortium (IMPC) and its NIH-funded Knockout Mouse Production and Phenotyping (KOMP^2^) component is producing a null allele mouse line for every protein coding gene in the mouse genome and phenotyping each null allele line to elucidate gene function in human biology and disease (International Mouse Knockout et al. 2007). During Phase I, IMPC/KOMP^2^ utilized a library of C57BL/6N ES cells harboring targeted null and sophisticated, flexible “knockout first” alleles to produce chimeric mice for germline transmission (Valenzuela et al. 2003, Skarnes et al. 2011, Coleman et al. 2015), to phenotype null allele mouse strains, and to bank cryopreserved sperm for distribution to the scientific community (Bradley et al. 2012). To date this effort has resulted in the phenotyping and distribution of thousands of null allele and conditional null allele lines (Ring et al. 2015). However, the need to generate and phenotype thousands more mutant mouse lines during Phase II funding has driven IMPC/KOMP^2^ to re-evaluate the use of targeted ES cell libraries, and adopt more efficient CRISPR/Cas9 gene editing approaches to supply their phenotyping pipelines with mice harboring critical exon deletion (null) alleles (Li et al. 2013a, Li et al. 2013b, Sung et al. 2014, Zuo et al. 2017). To maximize allele diversity available for distribution to the scientific community, developing CRISPR/Cas9 genome editing approaches amenable to high-throughput production of mice harboring conditional null alleles, in addition to null alleles, is of high priority to IMPC/KOMP^2^.

Here we report the results of a large-scale pilot study testing the feasibility of using CRISPR/Cas9-mediated HDR with pairs of sgRNAs and ssODNs to produce conditional null allele mice for the high-throughput IMPC/KOMP^2^ production pipelines. Our results demonstrate that CRISPR/Cas9-mediated HDR with pairs of ssODNs can successfully produce conditional null alleles at multiple loci across the genome. However, integration of loxP sites in *trans* and mutagenesis of loxP sequences impedes efficient conditional null allele production when using paired ssODNs. By contrast, our data suggest that CRISPR/Cas9-mediated HDR with pairs of sgRNAs a single lssODN harboring short homology arms and a loxP-flanked exon is more efficient than ssODNs at producing conditional null alleles and is therefore more appropriate for high-throughput conditional null allele mouse production. Importantly, both approaches present the opportunities of producing both null and conditional alleles when Cas9-induced DSBs are resolved by NHEJ rather than HDR. Thus, conditional alleles for the IMPC repository and null alleles for the phenotyping pipeline can be generated in the same microinjection. Together, our results indicate that lssODNs are likely to be the best, first choice for scaled conditional null allele production, and that ssODNs are a viable secondary option when conditional allele designs are not amenable to the lssODN approach.

## RESULTS

### Conditional allele design and genotyping schemes for CRISPR/Cas9-mediated HDR with ssODNs

To test whether CRISPR/Cas9-mediated HDR with pairs of sgRNAs and ssODNs can be used to efficiently and reliably produce conditional null alleles across the genome, we selected 30 genes to target for CRISPR/Cas9 genome editing. Genes selected for conditional alleles were either investigator-requested or sourced as previous failures of ES cell-based targeting by the IMPC. To capitalize on resources already available from the IMPC (www.mousephenotype.org), we utilized EUCOMM tm1a targeting vector designs (Bradley et al. 2012) already created for each gene to identify the critical exon or exons to flank with loxP sites. The following criteria were used to select critical exons to flank with loxP sites (“flox”). The 5’-most exon that: (1) is larger than 100 base pairs (bp) in size, (2) allows translation to initiate at the native start site, and (3) once removed from the genome by Cre/loxP recombination, induces a reading frameshift, a premature stop codon, and nonsense-mediated decay of all predicted coding transcripts. In some cases, multiple exons were selected for floxing as one single exon did not meet the 100 bp size requirement or induce a frameshift. All genes with MGI allele designs on the IMPC website have an accompanying GenBank file, which clearly annotates the critical exon or exons for targeting, listed as Ensembl exon IDs. Pairs of sgRNAs were identified to generate double-strand breaks within introns 5’ and 3’ to the critical exon(s). Selected target sequences were at least 100 bp 5’ or 3’ from the exon to be floxed (and any neighboring exons, when necessary) to minimize effects on splicing, ideally flanked by sequence without repetitive elements or GC content greater than 80% or less than 20% (Table S1).

For approximately two-thirds of the genes targeted for conditional allele production, we utilized ssODNs harboring conventional symmetrical homology arms 60 bases in length, not including the sgRNA sequence or the PAM site (Figure 1A, B), to generate a donor roughly 180 bases in length. To prevent unwanted mutagenesis of the correctly targeted allele, we modified the target site in the ssODN donor by placing the loxP sequence, along with a restriction enzyme recognition sequence, between bases 16 and 17 of the target sequence, 1 base away from the predicted cut site of Cas9 (Figure 1D). The sequence of the ssODN was complementary to the target strand. Richardson et al (2016) previously showed that ssODNs complementary to the non-target strand and with asymmetric homology arms (91 bases PAM-proximal, 36 bases PAM-distal) optimizes ssODN annealing to the DNA strand first released by Cas9 after DSBs are generated and increases the frequency of HDR events *in vitro*. Thereby, the remaining third of the genes targeted for CRISPR/Cas9-generated conditional alleles used asymmetrical homology arms to test if this approach increases HDR efficiency *in vivo* (Figure 1C). The location of the inserted loxP site was shifted within the sgRNA sequence, to be on the PAM-proximal arm, between bases 18 and 19 of the sgRNA sequence (Figure 1D) (Richardson et al. 2016).

**Figure 1 A-D.**
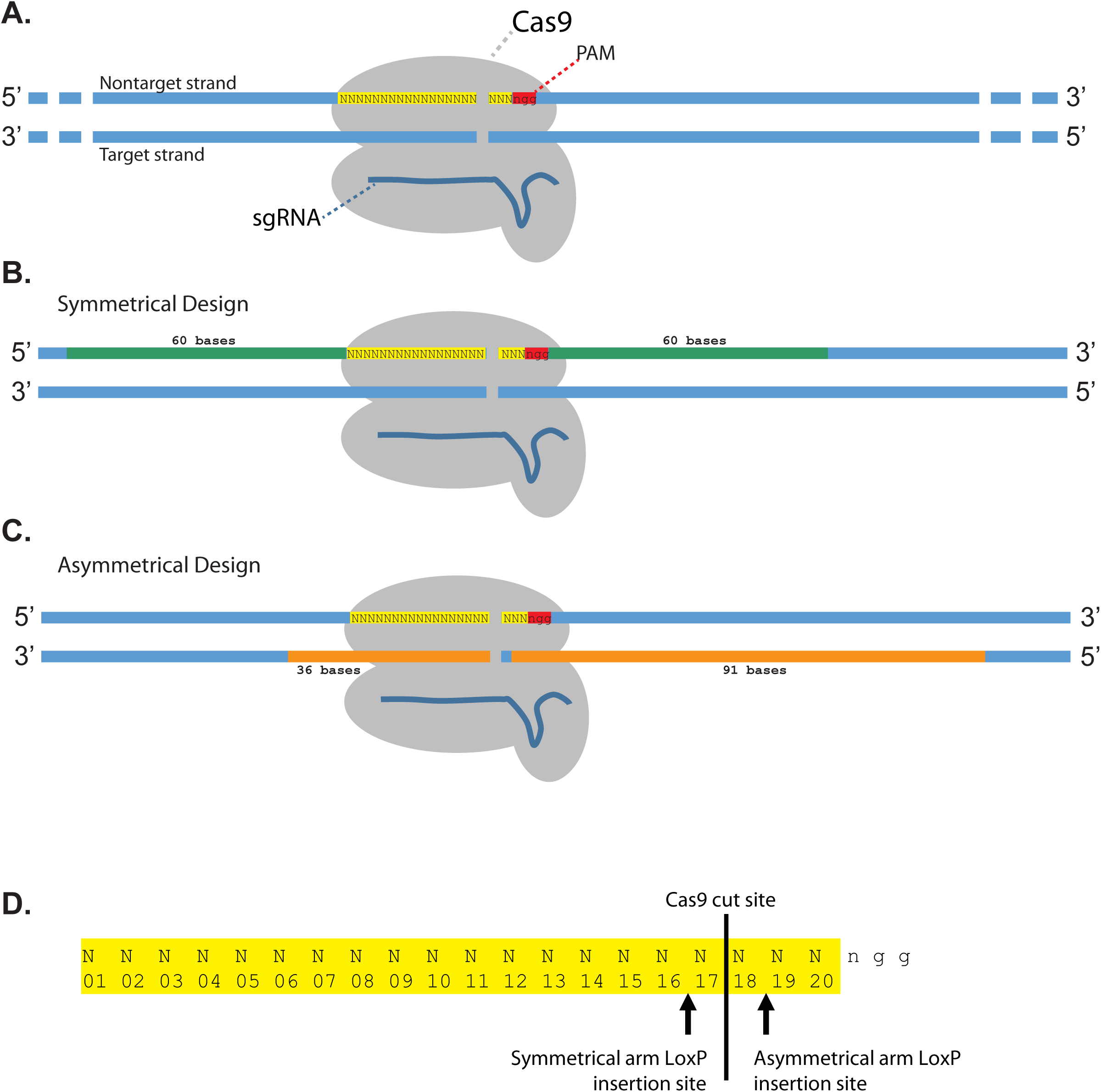
Designs for creating conditional alleles through CRISPR-mediated targeting with ssODN donor DNA. (A) Schematic for illustrating conditional targeting designs. Cas9 (gray) complexed with sgRNA (dark blue) binds to complementary DNA (blue) on the target strand after recognition of the PAM site (red). (B) Symmetrical design utilized two 60 bp homology arms, excluding the sgRNA target sequence and PAM site. The symmetrical ssODN donors were designed to be complementary to the target strand. (C) Asymmetrical design utilized a 36 bp PAM-distal and 91 bp PAM-proximal homology arms, as first described by Richardson et al (2016). The asymmetrical ssODN donors were designed to be complementary to the non-target strand. (D) Diagram illustrating the position of the loxP site insertion within the sgRNA target sequence in the ssODN donor. The loxP sequence was always inserted 1 base away from the Cas9 cut site, disrupting the sgRNA sequence in the ssODN donor and thereby preventing re-cutting by Cas9 after targeting.

Genotyping assays were designed to identify HDR-mediated insertion of each loxP site and NHEJ-mediated deletion of the critical exon between the two sgRNA target sites. To detect incorporation of each individual loxP site, PCR-based genotyping assays were designed to amplify at least 140 bp around the target sgRNA site for each loxP, with at least 1 primer outside the ssODN sequence (Figure 2A). Successful incorporation of the loxP site was identified by a 40 bp shift in the PCR product, representing the 34 bp loxP site and a 6 bp restriction enzyme site 5’ of the loxP sequence. To detect wild-type and critical exon deletion (null) alleles generated by NHEJ resolution of the two CRISPR/Cas9-generated double-strand breaks, a three primer PCR genotyping scheme was employed (Figure 2B). Both loxP PCR reactions and deletion/wild-type PCR reactions were performed on all F0 mice.

**Figure 2 A-C.**
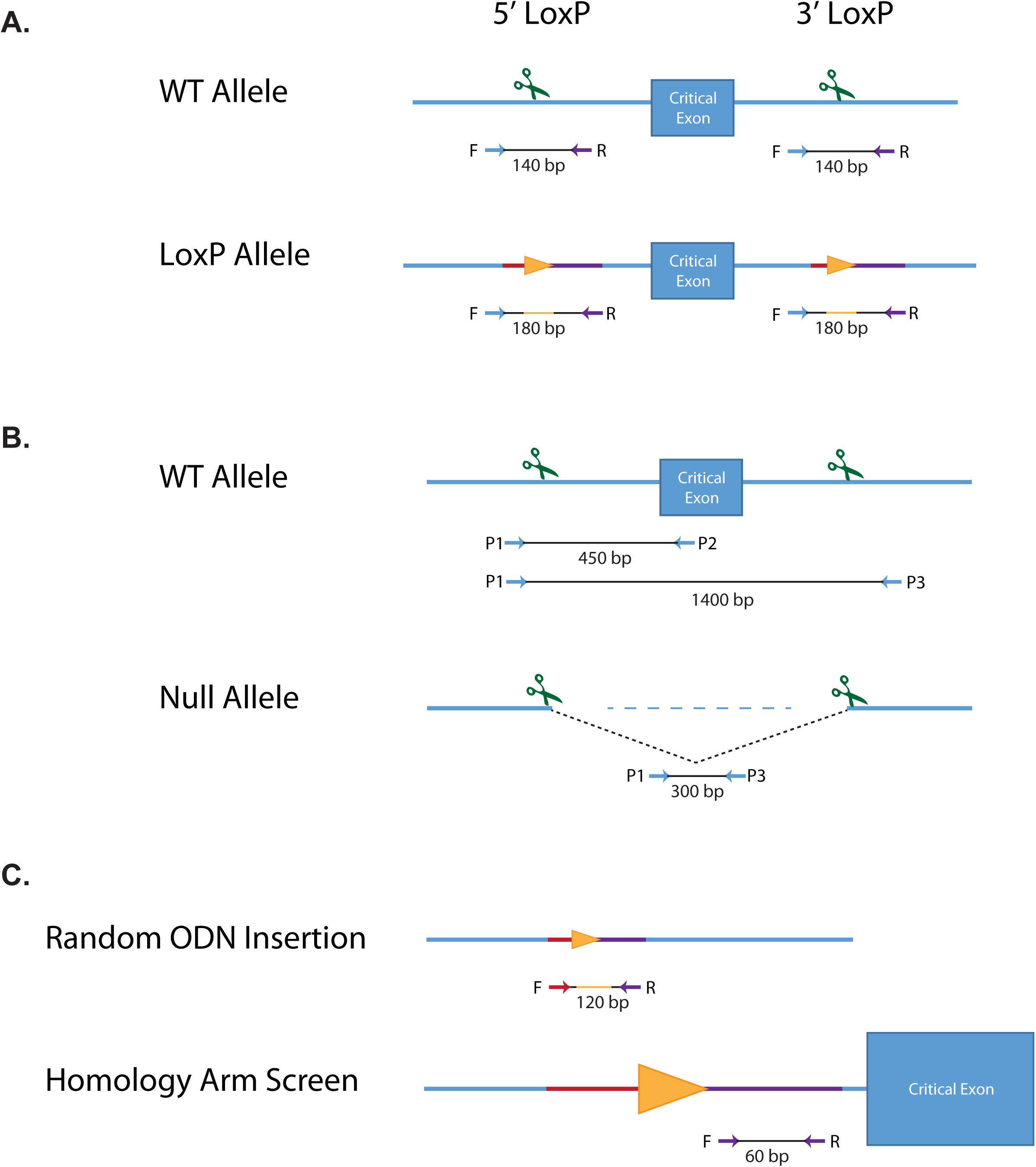
Genotyping for HDR and NHEJ alleles. Relative positions of the primers and approximate sizes of PCR products are listed below each allele. Scissors represent sgRNA sites. (A) Genotyping schemes for detecting loxP donor sequences. Orange triangles represent loxP sites, with representative homology sequence color coded on blue DNA strand. (B) Genotyping for NHEJ events utilizes a 3 primer system, with P1 being shared between P2 and P3. The P1+P3 primer pair may not always amplify a wild-type product, if the target sequences are too far apart. 1400 bp represents the average distance between loxP insertion sites. (C) Random ODN insertion PCR primers reside internal to homology arm sequence, and will amplify the expected size product if the ssODN donor has been incorporated elsewhere in the genome, away from the critical exon, in addition to the on-target locus. Primers for the homology arm screen were used in a SYBR-green quantitative PCR reaction with β-actin as a 2-copy control.

### CRISPR/Cas9-mediated HDR with ssODNs

To generate genome edited mice, pronuclear stage C57BL/6NJ embryos were microinjected with Cas9 mRNA, two sgRNAs, and two ssODNs into the cytoplasm. Approximately 200 embryos were microinjected per session per gene (Table S2). Litter sizes were noted for metrics of live-born pups based on the total number of zygotes injected. Microinjections and transfers resulted in fairly robust litters, with an average of 28 pups per attempt (14% of transferred embryos live-born).

Of the genes targeted with symmetrical homology arm donors, 15 of 22 genes had at least one putative founder mouse with both 5’ and 3’ loxP sites integrated into the genome; 21 of 22 genes had at least one live-born, genotyped (hereafter referred as F0) mouse with a null allele (Figure S1A and Table S2). In total, 7% of F0 animals from microinjection had both 5’ and 3’ loxP sites integrated; 18% harbored a null allele. Together, these results highlight the potential of generating two allele types with a single microinjection (Figure 3A).

**Figure 3 A-B.**
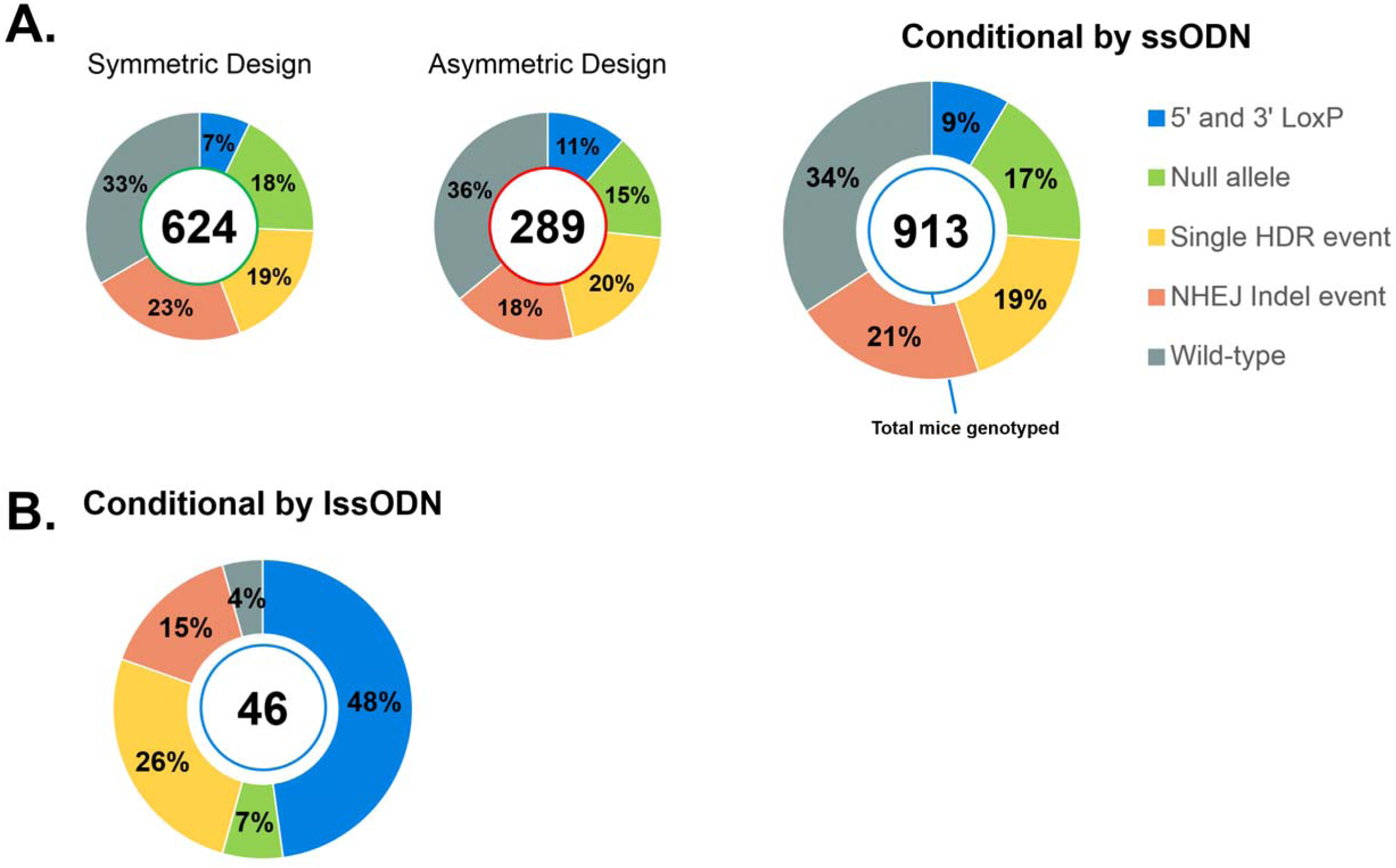
Conditional KO ssODN (A) and lssODN (B) targeting attempts. Each donut chart represents the summation of each allele type for all F0 mice genotyped, by ODN donor. In the center of the chart is the total number of F0 mice genotyped. Percentages of each allele type of the total number of mice genotyped are listed on each segment. 2 loxP: Includes animals genotyped for both 5’ and 3’ loxP sites, irrespective of the presence of any additional alleles (e.g., animals with 5’ loxP, 3’ loxP and a null allele detected); Null Allele: Includes animals genotyped for a null allele, which may also have a single HDR and/or NHEJ indel event; Single HDR: Includes animals genotyped for a single HDR event with or without additional indel events; NHEJ indel event: Animals in which only indel alleles were observed. (A) Summary of genotypes identified from ssODN targeting attempts, and symmetric and asymmetric homology arm designs. (B) Summary of genotypes identified from 4 lssODN targeting attempts.

Because previous reports suggested that using ssODNs with asymmetric homology arms improve HDR efficiency *in vitro* (Richardson et al. 2016), we next tested whether asymmetric homology arms increase HDR efficiency *in vivo*. Using the same sgRNAs, two loci (*Il1rl1* and *Eif2s2*) were targeted for conditional allele production using asymmetric and symmetric ssODNs designs (Figure S1B and Table S2). Targeting attempts with symmetric homology arm ssODNs successfully produced founders with 2 loxP sites at *Il1rl1* but not *Eif2s2*. Asymmetric homology arm ssODNs microinjections successfully produced animals with 2 loxP sites at *Eif2s2* but not *Il1rl1*. Microinjections using both asymmetric and symmetric ssODNs also produced founders with null alleles; therefore, failure to produce 2 loxP-containing animals was not the result of either sgRNA failing to induce DSBs. In summary, ssODNs with asymmetric homology arms appeared to not improve HDR efficiency at these two loci.

We attempted conditional allele generation at an additional 8 loci using asymmetrical homology arm ssODNs (Figure S1C), which resulted in 7 of 8 genes with at least one F0 mouse harboring both 5’ and 3’ loxP sites in the genome (11% of mice genotyped, Figure 3A). Furthermore, 15% of mice genotyped had a null allele, with at least one null allele founder in all 8 genes attempted by asymmetrical homology arm design. Direct comparison of putative founder production rates for 2 loxP sites between symmetric and asymmetric homology arm designs indicates a slight improvement from the asymmetric homology arm donor design (68% vs. 80%, respectively). Interestingly, there is a slight increase in the percentage of null alleles with symmetric homology arm designs, which could simply be attributed to the loci targeted or session effects during microinjection. Therefore, we did not detect any major differences between the two design strategies *in vivo*.

In summary for the 32 microinjections using pairs of ssODNs, 23 produced at least one F0 mouse with both 5’ and 3’ loxP sites integrated into the genome and 29 produced at least one F0 mouse with a null allele. Of all F0 mice, approximately 9% had 2 loxP sites integrated and 18% had a null allele (Figure 3A). For some targeting attempts, instances of both alleles were observed in a single mouse; 24% of the putative 2 loxP founder animals were identified with both 5’ and 3’ loxP sites and a null allele (Table S2).

Next, we were interested in whether there was any increased or decreased frequency of producing a putative conditional founder based on the genomic distance between the two targeted loxP sites. While more loxP insertions were attempted with smaller stretches of sequence between the two sgRNA target sites, there was no trend in loxP incorporation inefficiency with a specific distance (range of 250 to 4500 bp) (Figure 4A). Therefore, using paired ssODNs, critical exon(s) of varying sizes can floxed with reliable efficiency.

**Figure 4 A-B.**
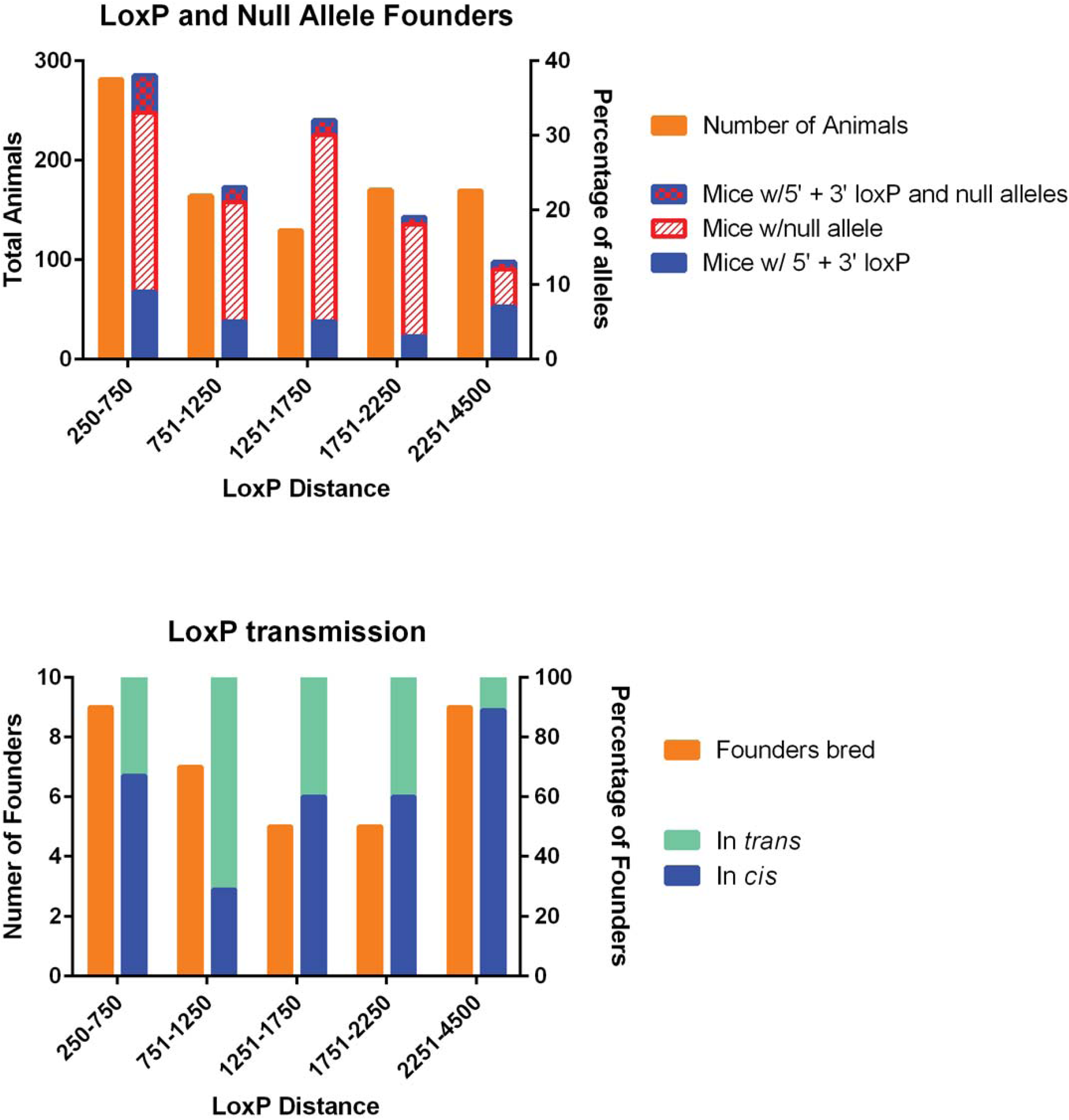
Frequency distributions of (A) projects with conditional, null, or both allele types, and (B) projects with loxP in *cis*, in *trans*, or both, binned by loxP distance. Project results derived from genotyping information gathered from F0 (A) and F1 (B) mice. LoxP distance calculated from genomic distance between Cas9 cut site of each sgRNA. X-axis labels indicate the median value of each bin. Left Y-axis depicts the number of projects attempted for each loxP distance bin, while the right Y-axis depicts the targeting or transmission percentage for all projects within each loxP distance bin.

### HDR efficiency with ssODNs is influenced by DSB efficiency

To determine whether HDR efficiency was influenced by the efficiency of sgRNA-guided, Cas9-mediated DSB production, we examined our loxP PCR genotyping results for PCR products indicative of NHEJ-generated indels at the sgRNA target sites (Figure 2A). The incidence of indel events at individual sgRNA target sites, the incidence of exon deletion events, which occur when double-strand breaks are generated at both the 5’ and 3’ sgRNA target sites, and the incidence of loxP insertion, which can only occur following a DSB event, from all F0 mice were then used to calculate the overall frequency of Cas9-generated DSBs at each target site. Cutting frequency was then compared to HDR frequency for all sgRNAs. Importantly, we detected a significant positive correlation between HDR and cutting frequency (Figure 5). Thus, sgRNA-guided, Cas9-mediated double-strand break production efficiency is a critical determinant of HDR efficiency.

**Figure 5.**
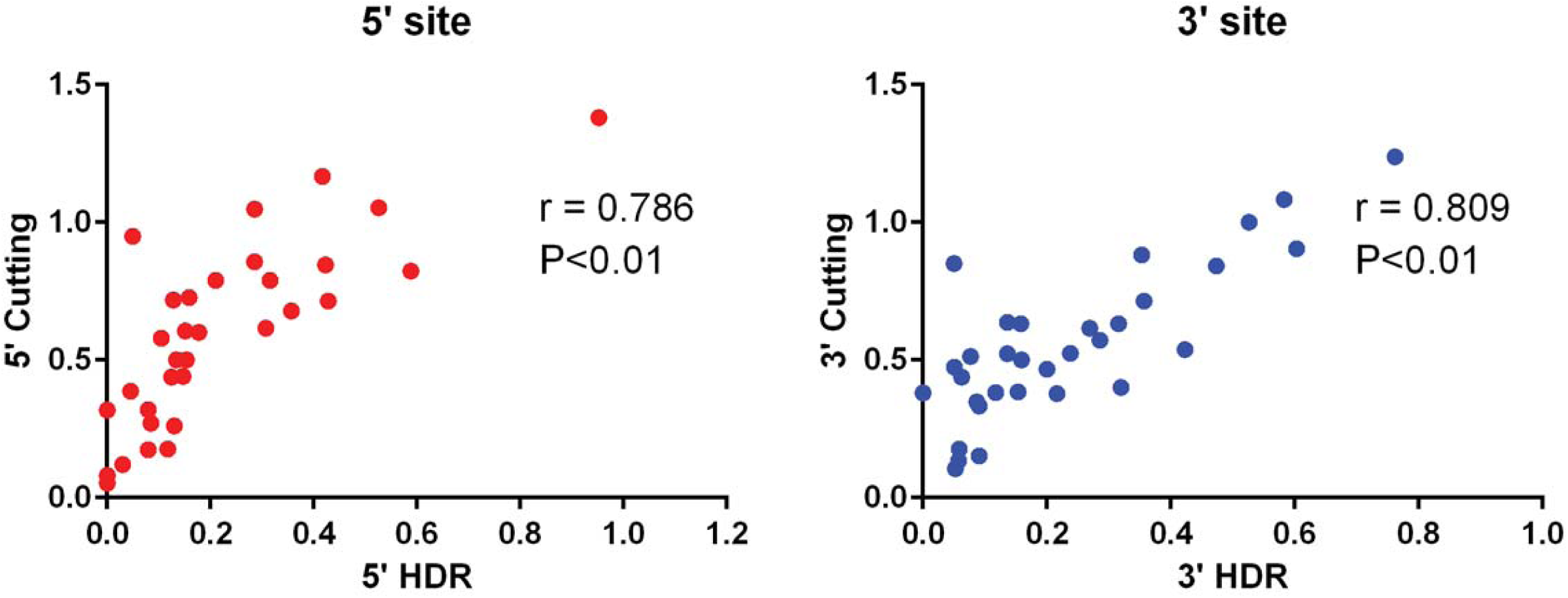
Pearson correlations for 5’ and 3’ loxP sites. Data plotted based on number of F0 mice with an HDR event (x-axis) vs. any evidence for a double-strand break generated at the respective sgRNA site, 5’ and 3’, such as a NHEJ indel, HDR event, or the formation of a null allele (y-axis).

### In *cis* vs in *trans* integration of loxP sites with ssODNs

One of the drawbacks of targeting loxP sites into the genome with two ssODNs is that the HDR events occur independently, with the potential for targeting to occur on separate chromosomes (in *trans)*. Thus, once founders harboring both 5’ and 3’ loxP sites were identified, we next tested for transmission of the loxP sites to the next generation as definitive proof that loxP sites were integrated on the same chromosome (in *cis*). For 22 lines having putative founders harboring both loxP sites, breeding identified at least one founder for 18 lines with the loxP sites in *cis* (81%, Table 1 and Table S2). On average, 63% of bred founders were found to transmit loxP sites in cis; 37% in trans. Interestingly, while there were few projects attempted with greater than 2.5 Kb between sgRNA target sites for loxP introduction, we did observe a greater frequency of loxP in *cis* integration when genomic distances between target sites was larger than 2250 bp (Figure 4B).

**Table 1.** Transmittance of 2 loxP sites to F1 generation.

| Donor | 2 loxP Loci Bred | loxP in cis | In cis, NOT mutated | Overall Success |
| --- | --- | --- | --- | --- |
| 2x ssODNs | 22 | 18 (81%)<sup>a</sup> | 14 (77%)<sup>b</sup> | **44%**<sup>c</sup> |
| symmetric | 15 | 14 (93%) | 10 (71%) | **45%** |
| asymmetric | 7 | 4 (57%) | 4 (100%) | **40%** |

<sup>a</sup>Percentage represents number of conditional alleles with loxP in cis over total 2 loxP loci bred; <sup>b</sup>Percentage represents number of loci with no mutations detected over all loci with loxP in cis. <sup>c</sup>Percentage represents number of loci with no mutations detected over all loci attempted.

Next, for the lines in which loxP sites were found to be in *cis*, Sanger sequencing was used to confirm the fidelity of the loxP sites in founder progeny. Of the putative founder mice selected for breeding from the 18 lines with loxP in *cis*, Sanger sequencing identified only 14 with at least one founder with both loxP sites intact (Table 1). The majority of mutated loxP sites observed by sequencing contained truncated sequences, although single base changes were also observed (Figure S2).

### Conditional allele design and genotyping schemes for CRISPR/Cas9-mediated HDR with lssODNs

As technology development has progressed since the adoption of CRISPR/Cas9 genome editing in mammals, methods for generating long (1000-2000 bp) single-stranded oligodeoxynucleotide donor molecules (lssODNs) have been published (Miura et al. 2015, Quadros et al. 2017). Miura et al., described a novel technique of using double-stranded DNA as a template for in vitro transcription of RNA from a T7 promoter sequence, and then reverse-transcribing the RNA to generate a single-stranded cDNA molecule. Quadros et al., demonstrated the feasibility of obtaining both conditional and reporter alleles at several loci using lssODN donor molecules in CRISPR-based targeting in mouse zygotes.

To test whether CRISPR/Cas9-mediated HDR with pairs of sgRNAs and a single, long ssODN (lssODN) could be used to efficiently and reliably produce conditional null alleles for high-throughput production, we targeted an additional 4 genes for CRISPR/Cas9 genome editing. We utilized resources already available from the IMPC for each gene to select the critical exon or exons to be “floxed”, and the genes to be targeted with this donor type had a critical region less than 1000 bp. Pairs of sgRNAs were identified to generate DSBs within introns 5’ and 3’ to the critical exon(s) as previously described for ssODNs (Table S1). For genes to be targeted for conditional alleles by lssODN (*Eif2s2, Mdh2, Megf11,* and *Cd44*), homology arms flanking the region to be floxed varied between 100-200 bp in length as needed to incorporate elements for synthesis of the lssODN (Miura et al. 2015, Quadros et al. 2017). For the 5’ homology arm, the length was determined starting at the 3’ end corresponding to base 17 in the 5’ sgRNA sequence (where the double-strand break would be initiated by Cas9) and terminating at the 5’ end with a trinucleotide stretch of guanines. A T7 promoter was added 5’ to the 2nd guanine, ensuring that the DNA donor template sequence contained 3 guanines at the 3’ end of the T7 promoter. For the 3’ homology arm, an anti-sense PCR primer was selected from a 200 bp window starting at base 17 in the 3’ sgRNA sequence, to be used to initiate reverse-transcription in the cDNA synthesis (Figure S3A).

Genotyping assays were designed to identify HDR-mediated insertion of each loxP site and NHEJ-mediated deletion of the critical exon between the two sgRNA target sites. To detect incorporation of each individual loxP site, PCR-based genotyping assays were designed to amplify approximately 350 bp around the target sgRNA site for each loxP, with at least 1 primer outside the lssODN homology arm (Supplemental Figure S3B). Successful incorporation of the loxP site was identified by a 34 bp shift in the PCR product. To detect critical exon deletion (null) alleles, the forward 5’ loxP primer and 3’ loxP reverse primer were used to amplify a deletion product. Both loxP and deletion PCR reactions were performed on all putative founder mice.

### CRISPR/Cas9-mediated HDR with lssODNs

To generate genome edited mice, pronuclear stage C57BL/6NJ embryos were microinjected with Cas9 mRNA, two sgRNAs, and a single lssODN into the cytoplasm. Approximately 200 embryos were microinjected per session per gene (Table S2). Microinjections and transfers resulted in notably smaller litters than microinjections with 2 ssODNs, with an average of 12 pups per attempt (8% of transferred embryos live-born).

Of the genes targeted with lssODNs, all had at least one F0 mouse with both 5’ and 3’ loxP sites integrated into the genome (Figure S1D). A much larger proportion of genotyped mice had both 5’ and 3’ loxP sites targeted, 48% of F0 mice from the lssODN microinjections, compared to 9% of F0 mice from ssODN microinjections. In comparison across donor types using the same sgRNAs for *Eif2s2*, putative founders were identified in 0% and 9% F0 mice with symmetric and asymmetric ssODNs, respectively, while the lssODN donor microinjection identified 67% of F0 mice with 2 loxP sites. Animals with null alleles occurred less frequently (7% of mice genotyped) when using lssODN in comparison to microinjections using pairs of ssODN (18% of mice genotyped). F0 conditional mice selected for breeding had loxP sites and floxed exons sequenced concurrently. Founder breeding from all four genes successfully transmitted the conditional allele, with both loxP in cis (Table S2). However, one of the bred founders had several single base mutations in the loxP sites.

Two of the four microinjections using lssODNs produced no animals with null alleles, which could be attributed to the loci attempted or session effects during the microinjection. Direct comparison of *Eif2s2* targeting across different donors produced null alleles in 16% and 18% F0 mice with symmetric and asymmetric ssODNs, respectively, while the lssODN donor microinjection produced no mice with a null allele (Figure S1B, D). A larger sample size would be needed to assess if there was a significant decrease in the ability to generate both conditional and null allele founders from a single microinjection with an lssODN.

### Off-target mutagenesis and random ODN insertions

Although the incidence of CRISPR/Cas9 off target mutagenesis appears to be significantly lower in mouse zygotes compared cell lines (Iyer et al. 2015, Bishop et al. 2016, Nakajima et al. 2016), off target mutagenesis is still possible, in particular for sgRNAs with off target sites harboring 1-2 mismatches (Iyer et al. 2015, Szafranski et al. 2017). Thus, to minimize the probability of off target events (Hsu et al. 2013), only sgRNAs predicted to have off-target sites with 3 mismatches or more were employed. To rule out off-target events in our conditional null allele lines, High Resolution Melt analysis of N1 animals derived from founders for 7 targeted genes were screened for off-target mutagenesis at the top 3 predicted off target sites for each sgRNA. N1 mice were selected for screening over founders due potential mosaicism in founders, which might obscure detection of an off-target mutagenesis event. For all off-target sites screened, we did not detect mutagenesis (see Table S3 for off-target locations).

The potential also exists for donors to randomly integrate into the genome. To screen for random ODN insertion (ROI), genotyping screens with primers internal to the donor ssODN or lssODN were designed to be run on all F0 mice from a subset of microinjections (Figure 2C and Table 2). The results of this genotyping were compared to the genotyping performed to detect the targeting of a loxP site, which uses a design with at least 1 primer outside the homology arms and detects only the on-target allele. Therefore, if an animal had a loxP site detected by the ROI screen, but not the on-target loxP genotyping screen, we consider this a random insertion event. F0 mice from 6 ssODN and 2 lssODN microinjections were screened for ROI events. Of the ssODN microinjections, 5 of 6 had at least 1 mouse with an ROI event (Table 2), with an observed frequency of 3-18% of mice affected. Both of the lssODN microinjections identified animals with an ROI event, with an observed frequency of 20-30% of mice affected. Therefore, random integration of single-stranded DNA donors does occur.

**Table 2.**
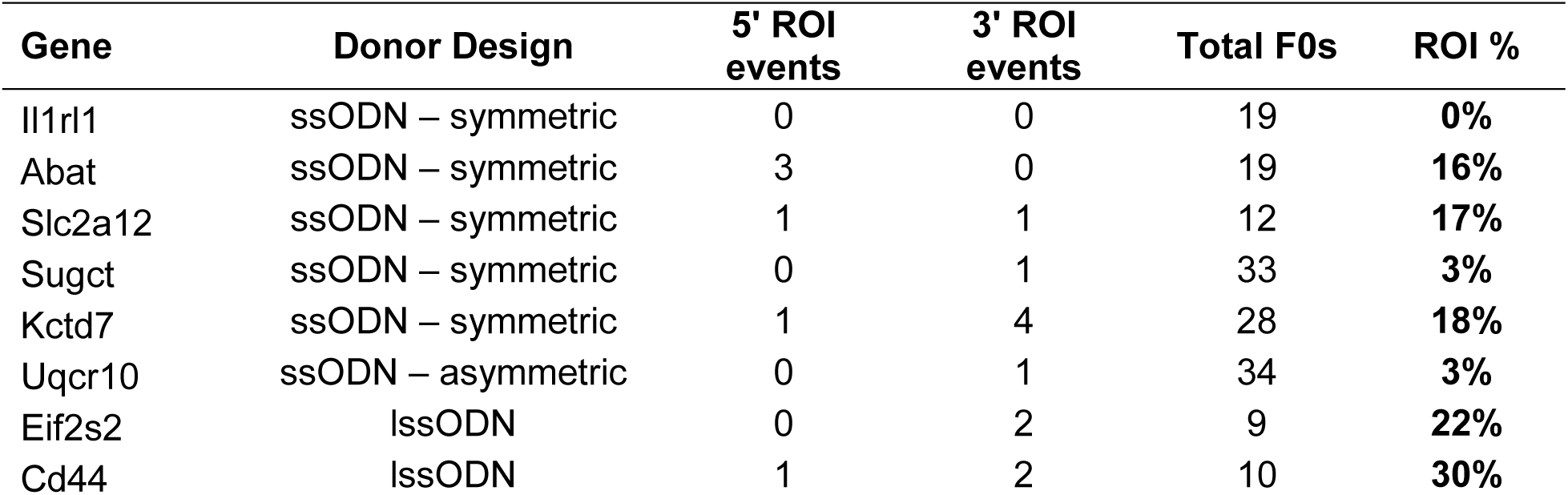
Founder screening for Random ODN Insertion

Because our initial ROI PCR screen could not identify ROI in animals harboring an on-target allele, we used alternative ROI assays. Quantitative SYBR Green real-time PCR using primers that amplify a portion of the homology arm was used to determine sequence copy number in ssODN-targeted N1 animals generated through founder backcrosses to wild-type C57BL/6NJ mice (Figure 2C). TaqMan copy number assays with probes internal to the critical exon were used to determine sequence copy numbers in lssODN targeted, N1 animals also produced through founder backcrosses to wild-type C57BL/6NJ mice. Conditional N1 offspring should have a homology arm or exon copy number of 2. Any N1 animal with more than 2 copies would be considered to have inherited both an on-target and ROI allele. Importantly, for both ssODNs and lssODNs, no increase in copy number was observed in any N1 animals screened (data not shown). Therefore, while random integration of single-stranded donor DNA is possible, the occurrence of both events occurring in the same animals appears to be rare.

## DISCUSSION

The effective generation of conditional alleles through the use of CRISPR/Cas9 has historically been challenging due to the inefficiencies of HDR in mouse zygotes. Plasmid donors have a relatively high cost and long lead time in production, and low frequency of targeting (Wang et al. 2013, Yang et al. 2013), which renders their use unsuitable for high-throughput production. Studies have attempted to shorten homology arms in plasmids from existing targeting vector libraries for their use as donors with CRISPR/Cas9 gene editing (Jung et al. 2017). However, any cost savings achieved through the use of existing plasmids is quickly lost in time needed to modify the donors prior to their use with CRISPR/Cas9. In comparison, ssODNs have proven to be simple to design, relatively cheap to produce, and an efficient donor molecule for CRISPR/Cas9-mediated HDR in mouse zygotes. While ssODN donors have been widely used to introduce sequence alterations at single sites (e.g. point mutations and epitope tags) (Wang et al. 2013, Wu et al. 2013, Yang et al. 2013, Bialk et al. 2016), their use for conditional allele production (two sites of HDR at the same locus) has been more limited in scale (Yang et al. 2013, Lee and Lloyd 2014, Bishop et al. 2016, Miano et al. 2016, Ma et al. 2017). More recently, lssODNs have proven to be an effective donor for CRISPR/Cas9-mediated insertion of floxed exons into the genome of mouse zygotes (Quadros et al. 2017), but again the scope of this experiment was limited. In the present study we have tested the use of CRISPR/Cas9-mediated HDR with ssODN pairs and single lssODNs to generate conditional null allele mice for high-throughput production. Based on our results and those of others who have employed ssODNs and lssODNs (Miura et al. 2015, Quadros et al. 2017), we propose that lssODN are the best donor template for reliable and efficient scale production of mouse lines harboring conditional alleles.

### Conditional allele production efficiency with ssODNs and lssODNs

When we compare conditional allele production efficiency between the two different single-stranded DNA donor types, lssODNs were substantially more efficient than ssODNs at producing F0 mice harboring intact and in *cis* loxP sites (48% vs 9%, respectively). The requirement for two independent HDR events when using ssODNs versus one HDR event when using lssODNs significantly contributed to differences in conditional allele production efficiency. Using pairs of sgRNAs and ssODNs, both sgRNAs need to efficiently facilitate Cas9-generated DSB production if HDR is to insert both loxP sites into the genome. Our data (Figure 5) clearly demonstrates that HDR efficiency is directly affected by the efficiency of DSB production. Thus, if one Cas9-sgRNA complex is suboptimal at generating a DSB, the rate of conditional allele production will be significantly reduced. If time and resources are permitting, pre-screening for Cas9-sgRNA complexes that are highly efficient at generating DSBs, and designing ssODNs around those target sites would most likely increase the probability of targeting both loxP sites into the genome of the same mouse. Although 2 sgRNAs were employed with each single lssODNs to induce sequence replacement by HDR, it is possible that only one Cas9-generated DSB at only 1 of the 2 sgRNA target sites was needed to facilitate HDR, thereby increasing overall HDR efficiency with lssODNs.

A second obstacle for conditional allele production with ssODNs is achieving two independent HDR events on the same allele. However, of the 35 putative conditional allele founders resulting from paired ssODN injections and backcrossed to generate N1 mice, 22 loxP sites transmitted in *cis* (63%) and 13 transmitted in *trans* (37%). Thus, there appears to be a preference for HDR events to occur on the same allele. Furthermore, when distance between sgRNA target sites was considered as a factor, in *cis* integration seemed to be highly preferred over in *trans* integration (89% vs 11% of bred founders, respectively) when the target sites were greater than 2250 bp apart. Thus, employing pairs of ssODNs for conditional allele production might be an efficient approach for loci that require loxP integration at distances over 2250 bp. Importantly, of the putative conditional founders that were produced using lssODN donors and were subsequently bred, all transmitted both loxP sites in *cis*. Thus, in *cis* integration of loxP sites is more reliable with lssODNs.

A recent study described a practice of generating conditional allele mice through microinjection or electroporation of one sgRNA and ssODN to target one loxP site into the genome at the one-cell embryo stage, followed by a subsequent microinjection or electroporation of a second sgRNA and ssODN to target the second loxP site into the genome of the same embryos at the two-cell stage (Horii et al. 2017). This approach decreased the potential of generating undesired interval deletion alleles in favor of the desired HDR (conditional) allele. Of the limited loci that the authors attempted, they observed decreased survival rates of zygotes translating into fewer live born animals, in addition to targeting rates marginally better than that reported herein using paired ssODN as donors in a single microinjection. Our high-throughput production to create conditional alleles would not be able to sustain the added costs or efforts required to target loxP sites sequentially.

### Null allele production when using ssODNs and lssODNs

From an IMPC/KOMP^2^ resource standpoint, it is highly advantageous to produce an exon deletion (null) allele and a conditional allele F0’s from the same microinjection. Thus, conditional allele mice for distribution to the scientific community and null allele animals for the phenotyping pipeline can be generated at the same time. Importantly, of the 831 F0 mice resulting from paired ssODN microinjections, 18% harbored a null allele. Furthermore, of the 9% of F0 mice harboring a putative conditional allele, 24% also harbored a null allele (Table S2). Animals that showed evidence of both NHEJ and HDR events were especially valuable if there was no evidence of the wild-type allele in the genotyping. It could be inferred that separate targeting events had occurred on opposite alleles and that the loxP sequences had been targeted in *cis*. Importantly, as the distance between sgRNA target sites increased, the number of mice harboring null alleles decreased, in particular when the distance between target sites exceeded 2250 bp (Figure 4A). It is likely that NHEJ repairs the two DSBs generated by the two sgRNAs independently as the distance between the target sites increases, resulting in two small indels rather than an interval deletion. Not surprisingly, the increase HDR efficiency observed when using lssODNs resulted in a decrease in F0 mice harboring a null allele (7%). In fact, only 2 of the 4 lssODN injections generated F0’s with a null allele. Thus, although lssODNs are more efficient at producing conditional allele founders, this comes at the expense of null allele founder generation.

### Mutagenized sequences at sites of HDR

Unfortunately, we did observe a proportion of successfully targeted loxP sites with mutagenesis when using paired ssODNs, which together with in *trans* integration, substantially reduced the efficiency of conditional allele founders. Two theories have been postulated to account for mutations observed. One hypothesis accounts for microhomology between a region of the donor DNA and the endogenous allele (Yoshimi et al. 2016), which may cause indel mutations due to microhomology-mediated end joining (MMEJ). Alternatively, synthesis errors when producing the ssODNs may become apparent as single copies are integrated into genomic DNA. A recent publication targeting loxP sites in the *Dock7* locus sequence-confirmed 1-to 2-bp deletions and a 1-bp substitution within the loxP site and/or surrounding sequence arising from incorrectly synthesized donor DNA (Bishop et al. 2016). Sequencing of N1 mice revealed single base deletions, which might be more attributable to synthesis errors, but large 10-12 base deletions (Figure S2) seem to suggest MMEJ. Using lssODNs, we also observed a founder with a successfully targeted floxed exon but with single base mutations within loxP sites. This alteration can most likely be attributed to a synthesis errors in the gBlock DNA template or acquisition of an error during the *in vitro* transcription or the cDNA synthesis steps of ssODN production. Thus, when using either ssODNs or lssODNs as DNA templates for conditional allele production, it is imperative to sequence across the site of HDR.

Importantly, although introduction of sequence errors in loxP sites (ssODNs or lssODNs) or floxed exons (lssODNs) are fatal flaws, errors in intronic sequences flanking loxP sites or exons can be tolerated if the base change(s) are predicted to not alter splicing or gene expression. Although altered intronic sequences are undesirable in the age of precision genome engineering, they have in the past been tolerated and proven to be mostly inconsequential when producing conditional null allele mice using traditional ES cell-based gene targeting. For example, ES cell clones harboring tm1a knockout first alleles and used by IMPC/KOMP^2^ to generate and phenotype thousands of knockout mouse lines have intronic microdeletions at sites where loxP sequences were introduced in the targeting vectors (Skarnes et al. 2011).

### Random insertion of ODNs

Importantly, our results indicate that both ssODNs and lssODNs can randomly integrate into the genome. However, of the founders harboring on-target conditional alleles and screened for ROI, we did not detect random integration. It is statistically unlikely that ROI and on-target HDR, two independent and somewhat rare events, will happen in the same animal. For ssODNs, of the 145 F0 animals analyzed for ROI, only 12 (8%) were found to have a ROI event (Table 2). Only 9% of F0 animals targeted with ssODNs harbored both a 5’ and 3’ loxP site (Figure 3). Thus, only 0.72% of F0s (8% x 9%) would be predicted to harbor a ROI and a putative conditional allele. However improbable that ROI and HDR events will occur in the same animal, our results demonstrate that it is important to screen for ROI in any mouse line produced through CRISPR-mediated HDR. Similar to random integration of transgenes, a ROI event could alter gene expression or function at the site of integration, and produce phenotypes not associated with the on-target modifications. Importantly, as long as ROI occurs on a separate chromosome from the on-target locus, breeding can be used to segregate the two alleles.

### ssODN homology arm symmetry

ssODN arm symmetry had varying effects on HDR efficiency, the rate of in *cis* integration, and mutagenesis at the site of HDR. Overall success, defined as projects with one founder harboring in *cis* loxP integration without sequence errors, was similar for symmetric and asymmetric ssODNs (45% vs 40% respectively, Table 1), but the success rates appear to be driven by different factors. Success at producing at least one founder with in *cis* integration of loxP sites was better for symmetric than asymmetric ssODNs (93% vs 57% of projects, respectively) while success at producing a founder with unaltered in *cis* loxP sites was better for asymmetric than symmetric ssODNs (100% vs 71% of projects with an in *cis* founder, respectively). These observed differences could be due to the loci selected for targeting. *Il1rl1* and *Eif2s2* were targeted using both symmetric and asymmetric ssODNs, but had opposite results: Symmetric ssODNs performed better for *Il1rl1* and asymmetric ssODNs worked better for *Eif2s2* (Figure S1B). However, aspects of the designs may have some influence on outcomes, in particular loxP mutagenesis. The asymmetric ssODN design employed (Richardson et al. 2016) used sequences complementary to the non-target strand and introduced novel sequences (i.e. loxP) on the PAM proximal side of the Cas9 cut site (Figure 1D). As previously reported, distal homology arms of asymmetric ssODNs appear to initiate strand invasion during synthesis-dependent strand annealing-mediated HDR (Richardson et al. 2016). Thus, loxP sequences in asymmetric ssODN might have been protected from potential mismatches with genomic DNA during repair. In comparison, loxP sites were introduced on the PAM distal side of the Cas9 cut site of target sequences when using symmetric ssODNs. Additional studies testing loci in parallel with each donor design would be necessary to verify that our observed differences in in *cis* integration and loxP mutagenesis are due to differences in homology arm symmetry and loxP placement relative to the Cas9 cut site.

### Considerations for using lssODNs

Importantly, although lssODNs harboring floxed exon sequences appear to more efficient than paired ssODNs harboring loxP sites at producing conditional alleles, there are several limiting factors to lssODNs that must be considered. In order to generate lssODNs via *in vitro* transcription and cDNA synthesis, a double-stranded DNA template is required. If the critical region exceeds the limits of synthetic DNA blocks currently commercially available, or if the sequence is unable to be synthesized as a double-stranded DNA template due to its complexity or repetitive nature, plasmid-cloned DNA can be used as a template. However, use of plasmids will increase time and cost of production. Furthermore, *in vitro* transcription and reverse transcription are error-prone and can introduce unwanted mutations in loxP sequences or exon sequences during lssODN production. Importantly, the recent availability of synthesized lssODNs (Megamers, IDT DNA) may address sequence error issues with lssODN production, but limitations on sequence length (up to 2000 bases) and synthesis of complex/repetitive sequences can still be problematic. With current methodologies, the use of paired ssODNs as DNA templates for CRISPR/Cas9-mediated HDR may be the only option for generating conditional alleles at some loci.

Our results demonstrate that CRISPR/Cas9-mediated HDR with lssODNs is currently the most efficient option for large scale production of mice harboring conditional null alleles. Furthermore, our results show that ssODNs, while significantly less efficient than lssODNs, can be a viable alternative to for conditional allele production when an lssODN donor is not feasible. While our efforts to date have focused on approaches for scaled production of mice harboring conditional null allele for IMPC/KOMP^2^, we will continue to test methods for generating larger and more complex lssODNs for scaled production of mice harboring knock-in or flexible alleles using CRISPR/Cas9 gene editing.

## MATERIALS AND METHODS

### Mouse strains

C57BL/6NJ and ICR mice were purchased from the Jackson Laboratory (Bar Harbor, ME) and maintained in the AAALAC-accredited animal facility at Baylor College of Medicine. All studies were reviewed and approved by the Institutional Animal Care and Use Committee of BCM and in accordance with National Institutes of Health guidelines for the Care and Use of Laboratory Animals.

### Conditional Allele design

Critical exon(s) to be floxed were identified using the targeting vector designs already created for each gene by the International Mouse Phenotyping Consortium (http://www.mousephenotype.org/). Single guide RNAs (sgRNAs) were selected using either the crispr.mit.edu or Wellcome Trust Sanger Institute (WTSI) Genome Editing websites; sgRNAs chosen had no fewer than 3 mismatches and were at least 100 bp proximal to the exon to be floxed and at least 100 bp from any neighboring exons. All sgRNA and off-target information can be found in links provided in Supplemental Tables 1 and 3, respectively. Symmetrical ssODN design used homology arms 60 bp in length, not including the sgRNA sequence or the PAM site. The loxP sequence was introduced between bases 16 and 17 of the sgRNA target site (predicted cut site of Cas9). The ssODN sequence was complementary to the target strand (Figure 1B). Asymmetrical ssODN design was attempted for 10 of the 30 genes targeted, which used asymmetric homology arms (91 bp PAM-proximal, 36 bp PAM-distal) and a ssODN sequence complementary to the non-target strand (Figure 1C). Novel restriction sites for *Xho*I, *Eco*RI, or *Bam*HI were placed immediately 5’ of the loxP integration site. LssODN design selected sgRNAs at least 50 bp 5’ or 3’ of the critical exons to be floxed. LoxP sites were introduced between bases 18 and 19 of the sgRNA target site and no restriction sites were included. T7 promoters were added to 5’ homology arms, at least 100 bp 5’ to the sgRNA site and 5’ to a trinucleotide stretch of guanines. The 3’ homology arm length was at least 100 bp, and terminated with a suitable primer for cDNA synthesis (Supplemental Figure 3A). Donor sequences are listed in Supplemental Table 1C; for the lssODN donors, the T7 promoter, loxP sites, and critical exon(s) are in uppercase, all other sequence in lowercase.

### Single stranded oligodeoxynucleotide donors

For ssODN donors, custom Ultramer^®^ DNA oligonucleotides were purchased from Integrated DNA Technologies (Coralville, IA). Synthesis of lssODN donors initiated with purchase of a DNA template from Integrated DNA Technologies (gBlock) (Supplemental Figure 3A). The DNA template was *in vitro* synthesized with mMESSAGE mMACHINE T7 Ultra transcription kit (ThermoFisher, cat. # AM1345) and purified using MEGAclear kit (ThermoFisher, cat. # AM1908) after DNase treatment, as described previously (Miura et al. 2015). The cDNAs were reverse transcribed from synthesized RNAs using SuperScriptIV Reverse Transcriptase (ThermoFisher, cat. # 18091050) using gene-specific primers (Supplemental Table 1), followed by RNase treatment. Unincorporated nucleotides and enzymes were removed from the cDNA by the Qiagen PCR purification kit (Cat. 28104) and the concentrations were measured using a NanoDrop™ 2000.

### sgRNA and Cas9 mRNA preparation

Single guide RNAs (sgRNAs) were synthesized using DNA templates for *in vitro* transcription. DNA templates were produced using overlapping DNA oligonucleotides in a high-fidelity PCR reaction (Bassett et al 2013). The PCR products were first purified using the QIAQuick PCR purification kit and used as a template for in vitro transcription of the sgRNA with the MEGAshort script T7 kit (ThermoFisher, AM1354). Following in vitro transcription, RNA was purified using the MEGAclear Transcription Clean-Up Kit (ThermoFisher AM1908). All samples were analyzed by Nanodrop to determine concentration and visualized using the Qiaxcel Advanced System using the RNA QC V2.0 kit to check the quality of RNA product before storage at -80°C. Cas9 mRNA was purchased from ThermoFisher (A25640). All sgRNAs were reanalyzed by Nanodrop prior to assembling the microinjection mixtures. Conditional allele attempts using ssODN donors consisted of Cas9 mRNA (100ng/μL), sgRNA (20 ng/μL, each), and 2 ssODNs (100ng/μL, each) in a final volume of 60 μL 1xPBS (RNAse-free). Conditional allele attempts using lssODN donors consisted of Cas9 mRNA (50ng/μL), sgRNA (10 ng/μL, each), and 1 lssODN (50ng/μL) in a final volume of 60 μL 1xPBS (RNAse-free). Sequences of donor templates used for loxP insertion are listed in the supplemental material (Supplemental Table 1).

### Microinjection of CRISPR/Cas9 reagents

C57BL/6NJ female mice, 24 to 32 days old, were injected with 5 IU/mouse of pregnant mare serum, followed 46.5 hr later with 5 IU/mouse of human chorionic gonadotropin. The females were then mated to C57BL/6NJ males, and fertilized oocytes were collected at 0.5 dpc. The BCM Genetically Engineered Mouse Core microinjected the sgRNA/Cas9/ssODN mixture into the cytoplasm of at least 200 pronuclear stage zygotes per gene attempted. Injected zygotes were transferred into pseudopregnant ICR females on the afternoon of the injection, approximately 25-32 zygotes per recipient female.

### Genotyping

Genomic DNA was isolated from ear punches from two week old pups using Sodium Hydroxide digestion followed by neutralization with Tris-HCl and dilution with nuclease-free water as previously described (Truett et al. 2000). Genomic DNA was isolated from tail clips of juvenile mice by proteinase K digestion, followed by isopropanol precipitation and resuspension in 1x Tris-EDTA. DNA was amplified by standard PCR using AmpliTaq Gold™ Fast PCR Master Mix (ThermoFisher, 4390939). To verify insertion of the loxP sites, primers were designed to PCR amplify a 140-180 base pair region around the sgRNA site, ensuring that at least one primer resided outside the homology arms of the ssODN (Figure 2A). To detect wild-type and null alleles generated by NHEJ, a three primer scheme was designed. Two primers bind outside the two sgRNA sites to PCR amplify deletion products. A third primer was designed to reside within the predicted deleted interval, to PCR amplify a product from the endogenous, wild-type allele (Figure 2B). All PCR products were visualized using the Qiaxcel Advanced System. Primer sequences used for genotyping are available upon request.

### DNA sequencing of cloned loxP sites in mice generated by CRISPR/Cas9

Genomic DNA was amplified as described above to visualize loxP insertions. The PCR products were cloned into competent cells using the pGEM Vector System according to the manufacturer’s protocol (Promega, A3600). Clones were screened by PCR for sequences containing a loxP site, and DNA Sanger sequenced by GENEWIZ (South Plainfield, NJ) for intact loxP sequences. Conditional alleles generated in founders from lssODN-targeted genes were sequenced by straight Sanger sequencing of purified PCR products generated from amplifying sequences around loxP sites and critical exon(s), and traces were aligned with SnapGene (GSL Biotech LLC) or deconvoluted using Sequencher (Gene Codes Corporation).

### Analysis of off-target Cas9 activity

The top three potential off-target sites for each sgRNA in the conditional targeting of *Il1rl1* (symmetric design targeting), *Abat, Kctd7, Slc2a12, Smc1a, Uqcr10,* and *Eif2s2* (lssODN targeting) were identified using the WTSI Genome Editing website. Flanking PCR primers designed to amplify 80-180 bp amplicons are listed in Supplemental Table 3 with the WGE ID, location, sequence, and number of mismatches from the original sgRNA. Off target mutagenesis was assessed by High Resolution MeltAnalysis using MeltDoctor HRM Master Mix (ThermoFisher, 4415440) on the QuantStudio 7 Flex Real-Time PCR System. At least 3 wild-type samples were analyzed concurrently with the test samples. For any sample with a HRM analysis result deviating from the wild-type sample, suggesting a mutagenesis event, PCR products were amplified with secondary sequencing primers and straight Sanger sequenced as described above.

### Analysis of random insertion of ssODNs

Genomic DNA was prepared from the F0 generation and standard PCR was performed using primers that flank the loxP sequence and resided within the homology arms of the ssODNs (Figure 2C). The results of the internal PCR genotyping were compared to the PCR genotyping used for on-target loxP insertion. To identify random insertions of ssODNs in lines established from founder animals harboring an on-target allele, primers were designed to PCR amplify sequences within the 5’ or 3’ homology arm of the ssODN for copy number counting in N1 animals. As described previously (D’Haene et al. 2010), DNA samples from ssODN-targeted N1 animals and wild-type C57BL/6NJ samples were amplified with the homology arm primers and *Actb* primers with Power SYBR™ Green PCR Master Mix (ThermoFisher, 4367659) for qPCR to serve as a copy number internal normalizing control. Homology arm copy number was determined relative to wild-type controls. DNA samples from lssODN-targeted N1 animals and wild-type control samples were analyzed by TaqMan^®^ Copy Number assays for *Eif2s2* and *Cd44* (ThermoFisher, Mm00054773_cn and Mm00049884_cn, respectively; Reference gene Tfrc 4458367).

## ACKNOWELDGEMENTS

We thank members of the International Phenotyping Consortium (IMPC) and Knockout Mouse Production and Phenotyping Project (KOMP^2^) CRISPR working group for consultation on allele design, reagent production, and microinjection conditions.

## FUNDING

This work was supported by NIH grants U42 OD011174 and UM1 HG006348 to the Knockout Mouse Production and Phenotyping (KOMP^2^) BaSH Consortium. Resources accessed through the BCM Mouse ES Cell and Genetically Engineered Mouse Cores were also supported by NIH grant P30 CA125123 to the Dan L. Duncan Cancer Center.

## REFERENCES

Andersson-Rolf, A., R. C. Mustata, A. Merenda, J. Kim, S. Perera, T. Grego, K. Andrews, K. Tremble, J. C. Silva, J. Fink, W. C. Skarnes and B. K. Koo (2017). “One-step generation of conditional and reversible gene knockouts.” Nat Methods 14(3): 287–289.

Bialk, P., B. Sansbury, N. Rivera-Torres, K. Bloh, D. Man and E. B. Kmiec (2016). “Analyses of point mutation repair and allelic heterogeneity generated by CRISPR/Cas9 and single-stranded DNA oligonucleotides.” Sci Rep 6: 32681.

Bishop, K. A., A. Harrington, E. Kouranova, E. J. Weinstein, C. J. Rosen, X. Cui and L. Liaw (2016). “CRISPR/Cas9-Mediated Insertion of loxP Sites in the Mouse Dock7 Gene Provides an Effective Alternative to Use of Targeted Embryonic Stem Cells.” G3 (Bethesda) 6(7): 2051–2061.

Boroviak, K., B. Doe, R. Banerjee, F. Yang and A. Bradley (2016). “Chromosome engineering in zygotes with CRISPR/Cas9.” Genesis 54(2): 78–85.

Bradley, A., K. Anastassiadis, A. Ayadi, J. F. Battey, C. Bell, M. C. Birling, J. Bottomley, S. D. Brown, A. Burger, C. J. Bult, W. Bushell, F. S. Collins, C. Desaintes, B. Doe, A. Economides, J. T. Eppig, R. H. Finnell, C. Fletcher, M. Fray, D. Frendewey, R. H. Friedel, F. G. Grosveld, J. Hansen, Y. Herault, G. Hicks, A. Horlein, R. Houghton, M. Hrabe de Angelis, D. Huylebroeck, V. Iyer, P. J. de Jong, J. A. Kadin, C. Kaloff, K. Kennedy, M. Koutsourakis, K. C. Lloyd, S. Marschall, J. Mason, C. McKerlie, M. P. McLeod, H. von Melchner, M. Moore, A. O. Mujica, A. Nagy, M. Nefedov, L. M. Nutter, G. Pavlovic, J. L. Peterson, J. Pollock, R. Ramirez-Solis, D. E. Rancourt, M. Raspa, J. E. Remacle, M. Ringwald, B. Rosen, N. Rosenthal, J. Rossant, P. Ruiz Noppinger, E. Ryder, J. Z. Schick, F. Schnutgen, P. Schofield, C. Seisenberger, M. Selloum, E. M. Simpson, W. C. Skarnes, D. Smedley, W. L. Stanford, A. F. Stewart, K. Stone, K. Swan, H. Tadepally, L. Teboul, G. P. Tocchini-Valentini, D. Valenzuela, A. P. West, K. Yamamura, Y. Yoshinaga and W. Wurst (2012). “The mammalian gene function resource: the International Knockout Mouse Consortium.” Mamm Genome 23(9-10): 580– 586.

Carroll, D. (2014). “Genome engineering with targetable nucleases.” Annu Rev Biochem 83: 409–439.

Chen, J., Y. Du, X. He, X. Huang and Y. S. Shi (2017). “A Convenient Cas9-based Conditional Knockout Strategy for Simultaneously Targeting Multiple Genes in Mouse.” Sci Rep 7(1): 517.

Chu, V. T., T. Weber, R. Graf, T. Sommermann, K. Petsch, U. Sack, P. Volchkov, K. Rajewsky and R. Kuhn (2016). “Efficient generation of Rosa26 knock-in mice using CRISPR/Cas9 in C57BL/6 zygotes.” BMC Biotechnol 16: 4.

Coleman, J. L., K. Brennan, T. Ngo, P. Balaji, R. M. Graham and N. J. Smith (2015). “Rapid Knockout and Reporter Mouse Line Generation and Breeding Colony Establishment Using EUCOMM Conditional-Ready Embryonic Stem Cells: A Case Study.” Front Endocrinol (Lausanne) 6: 105.

D’Haene, B., J. Vandesompele and J. Hellemans (2010). “Accurate and objective copy number profiling using real-time quantitative PCR.” Methods 50(4): 262–270.

Doudna, J. A. and E. Charpentier (2014). “Genome editing. The new frontier of genome engineering with CRISPR-Cas9.” Science 346(6213): 1258096.

Gaj, T., C. A. Gersbach and C. F. Barbas, 3rd (2013). “ZFN, TALEN, and CRISPR/Cas-based methods for genome engineering.” Trends Biotechnol 31(7): 397–405.

Horii, T., Y. Arai, M. Yamazaki, S. Morita, M. Kimura, M. Itoh, Y. Abe and I. Hatada (2014). “Validation of microinjection methods for generating knockout mice by CRISPR/Cas-mediated genome engineering.” Sci Rep 4: 4513.

Horii, T., S. Morita, M. Kimura, N. Terawaki, M. Shibutani and I. Hatada (2017). “Efficient generation of conditional knockout mice via sequential introduction of lox sites.” Sci Rep 7(1): 7891.

Hsu, P. D., D. A. Scott, J. A. Weinstein, F. A. Ran, S. Konermann, V. Agarwala, Y. Li, E. J. Fine, X. Wu, O. Shalem, T. J. Cradick, L. A. Marraffini, G. Bao and F. Zhang (2013). “DNA targeting specificity of RNA-guided Cas9 nucleases.” Nat Biotechnol 31(9): 827–832.

International Mouse Knockout, C., F. S. Collins, J. Rossant and W. Wurst (2007). “A mouse for all reasons.” Cell 128(1): 9–13.

Iyer, V., B. Shen, W. Zhang, A. Hodgkins, T. Keane, X. Huang and W. C. Skarnes (2015). “Off-target mutations are rare in Cas9-modified mice.” Nat Methods 12(6): 479.

Jung, C. J., J. Zhang, E. Trenchard, K. C. Lloyd, D. B. West, B. Rosen and P. J. de Jong (2017). “Efficient gene targeting in mouse zygotes mediated by CRISPR/Cas9-protein.” Transgenic Res 26(2): 263–277.

Lee, A. Y. and K. C. Lloyd (2014). “Conditional targeting of Ispd using paired Cas9 nickase and a single DNA template in mice.” FEBS Open Bio 4: 637–642.

Li, D., Z. Qiu, Y. Shao, Y. Chen, Y. Guan, M. Liu, Y. Li, N. Gao, L. Wang, X. Lu, Y. Zhao and M. Liu (2013a). “Heritable gene targeting in the mouse and rat using a CRISPR-Cas system.” Nat Biotechnol 31(8): 681– 683.

Li, W., F. Teng, T. Li and Q. Zhou (2013b). “Simultaneous generation and germline transmission of multiple gene mutations in rat using CRISPR-Cas systems.” Nat Biotechnol 31(8): 684–686.

Liu, E. T., E. Bolcun-Filas, D. S. Grass, C. Lutz, S. Murray, L. Shultz and N. Rosenthal (2017). “Of mice and CRISPR: The post-CRISPR future of the mouse as a model system for the human condition.” EMBO Rep 18(2): 187–193.

Ma, X., C. Chen, J. Veevers, X. Zhou, R. S. Ross, W. Feng and J. Chen (2017). “CRISPR/Cas9-mediated gene manipulation to create single-amino-acid-substituted and floxed mice with a cloning-free method.” Sci Rep 7: 42244.

Miano, J. M., Q. M. Zhu and C. J. Lowenstein (2016). “A CRISPR Path to Engineering New Genetic Mouse Models for Cardiovascular Research.” Arterioscler Thromb Vasc Biol 36(6): 1058–1075.

Miura, H., C. B. Gurumurthy, T. Sato, M. Sato and M. Ohtsuka (2015). “CRISPR/Cas9-based generation of knockdown mice by intronic insertion of artificial microRNA using longer single-stranded DNA.” Sci Rep 5: 12799.

Nakajima, K., A. A. Kazuno, J. Kelsoe, M. Nakanishi, T. Takumi and T. Kato (2016). “Exome sequencing in the knockin mice generated using the CRISPR/Cas system.” Sci Rep 6: 34703.

Quadros, R. M., H. Miura, D. W. Harms, H. Akatsuka, T. Sato, T. Aida, R. Redder, G. P. Richardson, Y. Inagaki, D. Sakai, S. M. Buckley, P. Seshacharyulu, S. K. Batra, M. A. Behlke, S. A. Zeiner, A. M. Jacobi, Y. Izu, W. B. Thoreson, L. D. Urness, S. L. Mansour, M. Ohtsuka and C. B. Gurumurthy (2017). “Easi-CRISPR: a robust method for one-step generation of mice carrying conditional and insertion alleles using long ssDNA donors and CRISPR ribonucleoproteins.” Genome Biol 18(1): 92.

Richardson, C. D., G. J. Ray, M. A. DeWitt, G. L. Curie and J. E. Corn (2016). “Enhancing homology-directed genome editing by catalytically active and inactive CRISPR-Cas9 using asymmetric donor DNA.” Nat Biotechnol 34(3): 339–344.

Ring, N., T. F. Meehan, A. Blake, J. Brown, C. K. Chen, N. Conte, A. Di Fenza, T. Fiegel, N. Horner, J. O. Jacobsen, N. Karp, T. Lawson, J. C. Mason, P. Matthews, H. Morgan, M. Relac, L. Santos, D. Smedley, D. Sneddon, A. Pengelly, I. Tudose, J. W. Warren, H. Westerberg, G. Yaikhom, H. Parkinson and A. M. Mallon (2015). “A mouse informatics platform for phenotypic and translational discovery.” Mamm Genome 26(9-10): 413–421.

Sander, J. D. and J. K. Joung (2014). “CRISPR-Cas systems for editing, regulating and targeting genomes.” Nat Biotechnol 32(4): 347–355.

Singh, P., J. C. Schimenti and E. Bolcun-Filas (2015). “A mouse geneticist’s practical guide to CRISPR applications.” Genetics 199(1): 1–15.

Skarnes, W. C., B. Rosen, A. P. West, M. Koutsourakis, W. Bushell, V. Iyer, A. O. Mujica, M. Thomas, J. Harrow, T. Cox, D. Jackson, J. Severin, P. Biggs, J. Fu, M. Nefedov, P. J. de Jong, A. F. Stewart and A. Bradley (2011). “A conditional knockout resource for the genome-wide study of mouse gene function.” Nature 474(7351): 337–342.

Sung, Y. H., J. M. Kim, H. T. Kim, J. Lee, J. Jeon, Y. Jin, J. H. Choi, Y. H. Ban, S. J. Ha, C. H. Kim, H. W. Lee and J. S. Kim (2014). “Highly efficient gene knockout in mice and zebrafish with RNA-guided endonucleases.” Genome Res 24(1): 125–131.

Szafranski, P., J. A. Karolak, D. Lanza, M. Gajecka, J. Heaney and P. Stankiewicz (2017). “CRISPR/Cas9-mediated deletion of lncRNA Gm26878 in the distant Foxf1 enhancer region.” Mamm Genome.

Truett, G. E., P. Heeger, R. L. Mynatt, A. A. Truett, J. A. Walker and M. L. Warman (2000). “Preparation of PCR-quality mouse genomic DNA with hot sodium hydroxide and tris (HotSHOT).” Biotechniques 29(1): 52–54.

Valenzuela, D. M., A. J. Murphy, D. Frendewey, N. W. Gale, A. N. Economides, W. Auerbach, W. T. Poueymirou, N. C. Adams, J. Rojas, J. Yasenchak, R. Chernomorsky, M. Boucher, A. L. Elsasser, L. Esau, J. Zheng, J. A. Griffiths, X. Wang, H. Su, Y. Xue, M. G. Dominguez, I. Noguera, R. Torres, L. E. Macdonald, A. F. Stewart, T. M. DeChiara and G. D. Yancopoulos (2003). “High-throughput engineering of the mouse genome coupled with high-resolution expression analysis.” Nat Biotechnol 21(6): 652–659.

Wang, H., H. Yang, C. S. Shivalila, M. M. Dawlaty, A. W. Cheng, F. Zhang and R. Jaenisch (2013). “One-step generation of mice carrying mutations in multiple genes by CRISPR/Cas-mediated genome engineering.” Cell 153(4): 910–918.

Wu, Y., D. Liang, Y. Wang, M. Bai, W. Tang, S. Bao, Z. Yan, D. Li and J. Li (2013). “Correction of a genetic disease in mouse via use of CRISPR-Cas9.” Cell Stem Cell 13(6): 659–662.

Yang, H., H. Wang, C. S. Shivalila, A. W. Cheng, L. Shi and R. Jaenisch (2013). “One-step generation of mice carrying reporter and conditional alleles by CRISPR/Cas-mediated genome engineering.” Cell 154(6): 1370– 1379.

Yoshimi, K., T. Kaneko, B. Voigt and T. Mashimo (2014). “Allele-specific genome editing and correction of disease-associated phenotypes in rats using the CRISPR-Cas platform.” Nat Commun 5: 4240.

Yoshimi, K., Y. Kunihiro, T. Kaneko, H. Nagahora, B. Voigt and T. Mashimo (2016). “ssODN-mediated knock-in with CRISPR-Cas for large genomic regions in zygotes.” Nat Commun 7: 10431.

Zuo, E., Y. J. Cai, K. Li, Y. Wei, B. A. Wang, Y. Sun, Z. Liu, J. Liu, X. Hu, W. Wei, X. Huo, L. Shi, C. Tang, D. Liang, Y. Wang, Y. H. Nie, C. C. Zhang, X. Yao, X. Wang, C. Zhou, W. Ying, Q. Wang, R. C. Chen, Q. Shen, G. L. Xu, J. Li, Q. Sun, Z. Q. Xiong and H. Yang (2017). “One-step generation of complete gene knockout mice and monkeys by CRISPR/Cas9-mediated gene editing with multiple sgRNAs.” Cell Res 27(7): 933–945.

